# Disposition of the chemosensory neurons in the neurotransmitter-release mutant *unc-13*

**DOI:** 10.1101/2024.07.21.604457

**Authors:** Eduard Bokman, Alon Zaslaver

**Affiliations:** Department of Genetics, Silberman Institute of Life Science, Edmond J. Safra Campus, The Hebrew University of Jerusalem, Israel

## Abstract

Secretion of neurotransmitters- and neuropeptides-containing vesicles is a regulated process orchestrated by multiple proteins. Of these, mutants, defective in the *unc-13* and *unc-31* genes, responsible for neurotransmitter and neuropeptide release, respectively, are routinely used to elucidate neural and circuitry functions. While these mutants result in severe functional deficits, their neuroanatomy is assumed to be intact. Here, using *C. elegans* as the model animal system, we find that the head sensory neurons show aberrant positional layout in neurotransmitter (*unc-13*), but not in neuropeptide (*unc-31*), release mutants. This finding suggests that synaptic activity may be important for proper cell migration during neurodevelopment and warrants considering possible anatomical defects when using *unc-13* neurotransmitter release mutants.

## Introduction

Animal genetic models offer powerful means to study gene-to-function relationships. One such model, the *C. elegans* nematode, is widely used to elucidate functional roles of individual genes and neurons (Corsi et al., 2015; Hobert, 2018). Moreover, owing to its fully-mapped 302-neuron connectome (Cook et al., 2019; White et al., 1986; Witvliet et al., 2021) and its compatibility with a myriad of genetic manipulations, *C. elegans* is instrumental to elucidate neuron-specific functions as well as network-wide principles that underlie various behavioral outputs (Randi and Leifer, 2020). In that respect, two mutants, the neurotransmitter (*unc-13)* and the neuropeptide (*unc-31)* release mutants continuously play pivotal roles (Richmond et al., 1999; Richmond and Broadie, 2002; Speese et al., 2007).

*Unc-13* is a highly conserved gene involved in docking the neurotransmitter-containing vesicles to the synaptic release sites, and subsequently priming them to be fusion-competent with the plasma membrane (Madison et al., 2005; Zhou et al., 2013; Hendi et al., 2019). Indeed, studies in worms, flies, and mice demonstrated that mutants, defective in *unc-13*, are impaired in neurotransmitter release (Aravamudan et al., 1999; Richmond et al., 1999; Varoqueaux et al., 2002). This impaired vesicle release was shown to affect transmission but not the layout and the general anatomy of the nervous system. For example, while *unc-13* defective *C. elegans* animals show severe uncoordinated movement, the architecture of the GABAergic motor neurons as well as their synaptic composition is intact (Richmond et al., 1999). Likewise, normal synaptic structures were observed in flies and mouse hippocampal neurons (Aravamudan et al., 1999; Varoqueaux et al., 2002).

The *unc-31* gene codes an essential component for the Ca-dependent exocytosis of dense-core vesicles (Speese et al., 2007; Hammarlund et al., 2008; Lin et al., 2010). These dense-core vesicles contain neuropeptides and monoamines that modulate presynaptic or postsynaptic functions via binding and activating G-protein coupled receptors. While both *unc-13* and *unc-31* participate in Ca-dependent exocytosis processes, in the *C. elegans* nervous system, they have parallel roles: *Unc-31* acts in dense-core vesicles while *unc-13* acts in synaptic vesicles.

In *C. elegans*, the head chemosensory system consists of 12 bilateral symmetric pairs of neurons (Bargmann, 2006). These neurons are located within the head amphids, laterally located sensilla that are open to the outside at the circumference of the worm’s mouth. Each neuron has a bipolar structure: anteriorly, it extends a neurite to the tip of the animal’s nose, while the other branch is an axon-like extension forms in- and out-going synapses with other neurons within a dense nerve ring. These neurons sense chemicals in the environment and play prominent roles in many physiological and behavioral outputs, including chemotaxis, mechanosensation, osmotaxis, thermotaxis and more (Bargmann, 2006; Bokman et al., 2024; Goodman et al., 2018; Goodman and Sengupta, 2019; Zaslaver et al., 2015).

Herein, we analyzed the anatomical position of the amphid chemosensory neurons in both *unc-13* and *unc-31* mutants. We find that while the anatomical layout of the neuropeptide release mutant *unc-31* is indistinguishable from that of WT animals, the cell bodies in the *unc-13* mutants are disarranged. This finding challenges current held assumptions that *unc-13* mutants are impaired in vesicle release only. Moreover, it suggests that intact neurotransmitter release may play important roles in shaping and organizing the anatomical layout the nervous system.

## Results and Discussion

To study the organizational layout of the amphid chemosensory neurons, we imaged a transgenic strain in which all amphid neurons express a florescent marker (**Fig 1A-C**). For imaging, we used a confocal microscope capturing z-stack images at a resolution of 0.6 µm apart, sufficient for cellular resolution detection of individual neurons. Using our previously developed image analysis pipeline (Iwanir et al., 2019; Pritz et al., 2023), we segmented and identified individual neurons based on nuclear expression of mCherry. Notably, mCherry nuclear localization is crucial as it provides the cellular resolution required for accurate identification of individual neurons. The result of this anatomical analysis generates the 3D organization of the cells nuclei **(Fig. 1D)**.

**Figure 1.**
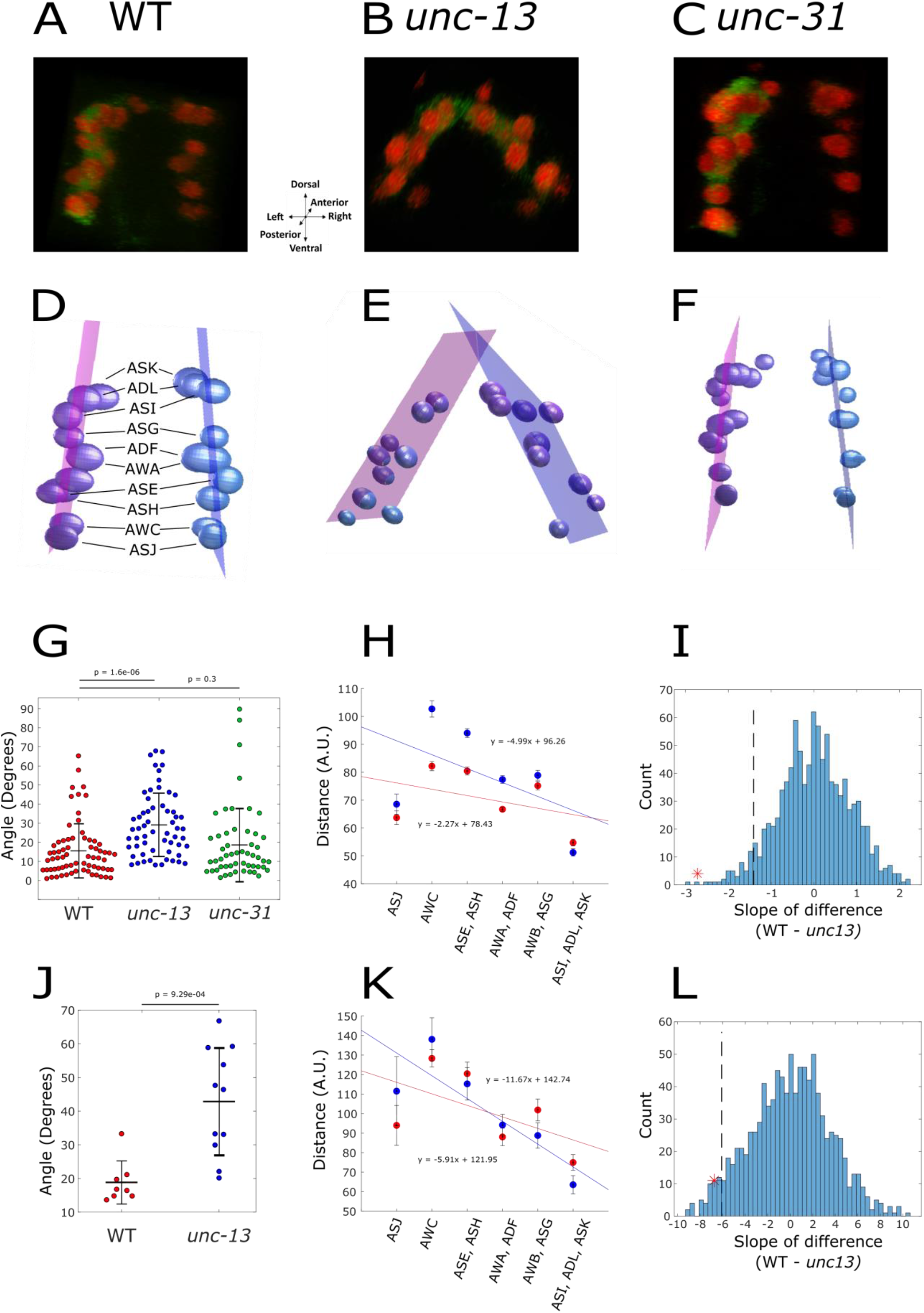
The anatomical layout of the chemosensory neurons is aberrant in *unc-13*, but not in *unc-31*, mutant animals. **(A-C) 3D imaging of the amphid chemosensory neurons in** (A) WT, (B) *unc-13* and (C) *unc-31* mutant animals. This image is produced by stacking together the individual Z-plane images. The imaged strain expresses GCaMP in the cytoplasm and mCherry in the nuclei of all amphid neurons (*osm-6*::GCaMP, *osm-6*::mCherry-NLS (Iwanir et al., 2019)). **(D-F)** Spatial localization of the nuclei of the amphid chemosensory neurons in (D) WT, (E) *unc-13* and (F) *unc-31* animals. An in-house image analysis pipeline segments, identifies, and labels the individual chemosensory neurons from both lateral sides of the amphid (Iwanir et al., 2019; Pritz et al., 2023). A best fit plane encompassing all the neurons in each side of the amphid was constructed. **(G-I)** The angle that two planes of the sensory neurons form at the dorsal side of the amphid (G), and the distance between the corresponding two bilateral neurons together with the regression lines (WT, red; *unc-13*, blue, panel H). A bootstrap analysis which calculates the slope for randomly chosen WT and *unc-13* indicated that the observed difference between the strains is statistically significant (p< 0.001, see methods). Dashed line represents the 5% percentile of the bootstrap estimated slope difference. Red asterisk denotes the observed slope difference. These measurements are from worms inserted into a microfluidic device. **(J-L)** Animal were placed on an agar pad and imaged. (J) The angle that the two planes of the sensory neurons form at the dorsal side of the amphid. (K) The distance between the corresponding two bilateral neurons together with the regression lines (WT, red; *unc-13*, blue, panel H). (L) Bootstrap analysis (as in panel I) to indicate the significance (p=0.029).

We next compared neural spatial organization of WT animals to that of *unc-31* and *unc-13* mutant strains **(Fig 1D-F)**. The neural layout of the *unc-31* mutants resembled that of WT animals, where the neurons within each of the two amphid halves were positioned in parallel. In contrast, the layout of the *unc-13* mutants showed a stark difference where the two amphid halves were tilted such that they are intersected at the dorsal part.

To quantitate this anatomical difference, and to provide a statistical measure for the significance of the aberrant organization, we measured the angle formed between the two bilateral sides of the amphid neurons. First, we constructed best-fit planes for the right and the left halves of the amphid (**Fig. 1D-F**). Each plane crosses the neurons positioned in one lateral side such that the summed distances of the nuclei from the plane is minimal (Methods). Indeed, these planes are almost in a perfect parallel position in WT and *unc-31* mutant animals, and the average angle between the planes are 15.51±14.18 degrees and 18.50±19.18, respectively **(Fig 1D,F,G)**. In contrast, in *unc-13* mutant strain, the two halves of the amphid intersect one another at the dorsal side forming a significantly larger angle between the two planes of 29.09±16.66.

A complementing approach to quantify neural displacement observed in the *unc-13* mutant is to calculate the distance between the corresponding bilateral right and the left types of neurons. We find that while the average distance between individual bilateral neurons is similar in WT and *unc-31* mutant animals, this distance is significantly higher in *unc-13* mutants (**Fig. 1H**). Specifically, the increased distance is observed between the internal neurons positioned along the dorso-ventral axis (namely, AWC, AWC, ASH, ASE, and ADF), whereas the distance between the five dorsal bilateral neurons (ADL, ASK and ASI, AWB, and ASG) and the most ventral bilateral neurons (ASJ) is intact and resembles the distance found in WT animals. To provide a quantitative measure for the greater distancing in the *unc-13* mutant, we calculated the regression line denoting the distance between the bilateral neurons in the anterior-posterior axis **(Fig 1H)**. We find that the slope of the *unc-13* mutant is two-fold higher than that of the WT, a statistically significant difference confirmed using a bootstrap analysis (p< 0.001, **Fig 1I**). Together, these results indicate that the abnormal neural position is mainly due to the greater distance between the bilateral neurons positioned in the inner part of the anterior-posterior axis of the animal.

The imaging procedure detailed above was performed on animals restrained in a microfluidic device (Chronis et al., 2007). This approach allows measuring accurate neural position by imaging multiple planes while the animals are held in place. However, inserting the worms into a tight restraining channel squeezes and pressurizes them. Consequently, it could be that the observed neural displacement of *unc-13* mutants is due to a reduced resistance to external mechanical forces, rather than neuroanatomical-related defects.

To distinguish between the two possibilities, we reimaged the WT and the *unc-13* mutants without restraining them in a microfluidic device. For this, worms were anesthetized and laid on an agar pad, such that they did not experience any external force (Methods). Comparing the angle between the two bilateral planes showed a significantly larger angle in the *unc-13* mutants **(Fig 3J)**, thus corroborating the findings using the microfluidic chips. In fact, while the angle of WT animals was very similar (18.84°±6.33), the angle of the *unc-13* mutant was larger than the one observed when using microfluidic devices (42.71°±15.79 vs 29.09°±-16.66). Thus, the difference in the angles is even more pronounced when not squeezing the animals inside the narrow microfluidic channels. Measuring the distance between the bilateral neurons showed that the in *unc-13*, the sensory neurons in middle of the anterior-posterior axis are further apart than the neurons at the edge on this axis, similarly to what was observed when imaging the neurons in the microfluidic device (**Fig 1J**). Likewise, the slope of the regression line between the neurons was two-fold higher than in the *unc-13* mutant (p=0.029, **Fig 1K-L**).

Altogether, here we show that a mutation in the *unc-13* gene, but not *unc-31*, leads to positional displacement of the chemosensory neurons cell bodies. This raises the interesting possibility that intact synaptic activity may be necessary for proper cell migration during development of the nervous system. While synaptic organization of the amphid neurons was not studied herein, previous studies in *C. elegans*, using the same *unc-13* mutant allele (*S69)*, found no anatomical changes in the GABAergic neuromuscular junctions, where the average interval between synapses and the distribution of the postsynaptic GABA receptors was indistinguishable from that of WT animals (Richmond et al., 1999). Similarly, while the priming and secretion of neurotransmitter vesicles were severely impaired, normal synaptic structures were observed in mutant flies, as well in mouse hippocampal neurons (Aravamudan et al., 1999; Varoqueaux et al., 2002). Thus, while dysfunctional synaptic activity may not affect proper synaptic formation, it may affect neurodevelopmental-related migratory processes of the cell bodies. Furthermore, these findings imply that when using *unc-13* mutants, in addition to neurotransmission impairment, one may also need to consider possible neuroanatomical defects.

## Materials and Methods

### Strains

ZAS280 *In*[*osm-6*::GCaMP3, *osm-6*::ceNLS-mCherry-2xSV40NLS] (Iwanir et al., 2019).

ZAS325 is a cross between ZAS280 and *unc-31*(e928) (Pritz et al., 2023).

ZAS371 is a cross between ZAS280 and *unc-13*(s69).

### Worm cultivation

All strains were grown on NGM plates pre-seeded with OP 50 and kept at 20 °C. Age synchronization was performed by bleaching. All experiments were done on young-adult animals, 3-days post bleaching.

### Imaging the chemosensory system

For microfluidic-based imaging, individual worms were inserted into a microfluidic chamber (Chronis et al., 2007) where they were partially anesthetized with 10 mM levamisole and left to habituate for 10 minutes (Bokman et al., 2024; Itskovits et al., 2018; Ruach et al., 2022). For the agar pad experiments a small think chuck of NGM agar was placed onto a slide. A ∼15µl drop of 10 mM levamisole was pipetted onto the agar, and small groups of 5-8 worms were picked into the drop, gently covered with a coverslip, and left to habituate for 10 minutes.

Imaging was performed on a Nikon A1R+ confocal laser scanning microscope with a water immersion x40 (1.15NA) objective Z-axis intervals of 0.5-0.8 µm. The system was controlled by the Nikon NIS-elements software.

### Neuron identification

Neural nuclei were segmented based on the nuclear mCherry signal using an algorithm developed by (Toyoshima et al., 2016). Neuronal identities were determined visually based on anatomical positions. Neurons that could not be unambiguously identified were removed from analysis.

### Statistical analysis

The angles between the two sides of the amphid were determined by fitting a plane to the 3-dimentional positions of the nuclei centroids on each side, and calculating the angle at which the planes intersect. Worms with fewer than 4 identified neuron pairs were excluded. P-values were obtained using a two-sample t-test.

For the distances within the bilateral neuron pairs, the neuron classes were assigned ranks based on their typical relative position along the dorso-ventral axis. Neuron that are close together on this axis were given the same rank. From ventral to dorsal the ranks are: ASJ = 1, AWC = 2, (ASE, ASH) = 3, (AWA, ADF) = 4, (AWB, ASG) = 5, (ASI, ADL, ASK) = 6. Distances between the left and right sides of each neuron pair were calculated, and a regression line was fit to the mean distances of the ranks. P-values were obtained using a permutation test. All distances within each rank were shuffled and randomly assigned to WT or mutant. The mean mutant distance per rank was then subtracted from the mean WT distance, and a regression line was fit to the result. The observed value of the slope of this regression line was compared to the slopes of 1000 random permutations.

## References

Aravamudan, B., Fergestad, T., Davis, W.S., Rodesch, C.K., Broadie, K., 1999. Drosophila Unc-13 is essential for synaptic transmission. Nat. Neurosci. 2, 965–971. 10.1038/14764

Bargmann, C., 2006. Chemosensation in C. elegans. WormBook. 10.1895/wormbook.1.123.1

Bokman, E., Pritz, C.O., Ruach, R., Itskovits, E., Sharvit, H., Zaslaver, A., 2024. Intricate response dynamics enhances stimulus discrimination in the resource-limited C. elegans chemosensory system. 10.1101/2024.01.14.575365

Chronis, N., Zimmer, M., Bargmann, C.I., 2007. Microfluidics for in vivo imaging of neuronal and behavioral activity in Caenorhabditis elegans. Nat. Methods 4, 727–731. 10.1038/nmeth1075

Cook, S.J., Jarrell, T.A., Brittin, C.A., Wang, Y., Bloniarz, A.E., Yakovlev, M.A., Nguyen, K.C.Q., Tang, L.T.-H., Bayer, E.A., Duerr, J.S., Bülow, H.E., Hobert, O., Hall, D.H., Emmons, S.W., 2019. Whole-animal connectomes of both Caenorhabditis elegans sexes. Nature 571, 63–71. 10.1038/s41586-019-1352-7

Corsi, A.K., Wightman, B., Chalfie, M., 2015. A Transparent Window into Biology: A Primer on Caenorhabditis elegans. Genetics 200, 387–407. 10.1534/genetics.115.176099

Goodman, M.B., Klein, M., Lasse, S., Luo, L., Mori, I., Samuel, A., Sengupta, P., Wang, D., 2018. Thermotaxis navigation behavior, in: WormBook: The Online Review of C. Elegans Biology [Internet]. WormBook.

Goodman, M.B., Sengupta, P., 2019. How Caenorhabditis elegans Senses Mechanical Stress, Temperature, and Other Physical Stimuli. Genetics 212, 25–51. 10.1534/genetics.118.300241

Hammarlund, M., Watanabe, S., Schuske, K., Jorgensen, E.M., 2008. CAPS and syntaxin dock dense core vesicles to the plasma membrane in neurons. J. Cell Biol. 180, 483–491. 10.1083/jcb.200708018

Hendi, A., Kurashina, M., Mizumoto, K., 2019. Intrinsic and extrinsic mechanisms of synapse formation and specificity in C. elegans. Cell. Mol. Life Sci. 76, 2719–2738. 10.1007/s00018-019-03109-1

Hobert, O., 2018. The neuronal genome of Caenorhabditis elegans, in: WormBook: The Online Review of C. Elegans Biology [Internet]. WormBook.

Itskovits, E., Ruach, R., Kazakov, A., Zaslaver, A., 2018. Concerted pulsatile and graded neural dynamics enables efficient chemotaxis in C. elegans. Nat. Commun. 9, 2866. 10.1038/s41467-018-05151-2

Iwanir, S., Ruach, R., Itskovits, E., Pritz, C.O., Bokman, E., Zaslaver, A., 2019. Irrational behavior in C. elegans arises from asymmetric modulatory effects within single sensory neurons. Nat. Commun. 10, 3202. 10.1038/s41467-019-11163-3

Lin, X.-G., Ming, M., Chen, M.-R., Niu, W.-P., Zhang, Y.-D., Liu, B., Jiu, Y.-M., Yu, J.-W., Xu, T., Wu, Z.-X., 2010. UNC-31/CAPS docks and primes dense core vesicles in C. elegans neurons. Biochem. Biophys. Res. Commun. 397, 526–531. 10.1016/j.bbrc.2010.05.148

Madison, J.M., Nurrish, S., Kaplan, J.M., 2005. UNC-13 interaction with syntaxin is required for synaptic transmission. Curr. Biol. CB 15, 2236–2242. 10.1016/j.cub.2005.10.049

Pritz, C., Itskovits, E., Bokman, E., Ruach, R., Gritsenko, V., Nelken, T., Menasherof, M., Azulay, A., Zaslaver, A., 2023. Principles for coding associative memories in a compact neural network. eLife 12, e74434. 10.7554/eLife.74434

Randi, F., Leifer, A.M., 2020. Measuring and modeling whole-brain neural dynamics in Caenorhabditis elegans. Curr. Opin. Neurobiol., Whole-brain interactions between neural circuits 65, 167–175. 10.1016/j.conb.2020.11.001

Richmond, J.E., Broadie, K.S., 2002. The synaptic vesicle cycle: exocytosis and endocytosis in Drosophila and C. elegans. Curr. Opin. Neurobiol. 12, 499–507. 10.1016/s0959-4388(02)00360-4

Richmond, J.E., Davis, W.S., Jorgensen, E.M., 1999. UNC-13 is required for synaptic vesicle fusion in C. elegans. Nat. Neurosci. 2, 959–964. 10.1038/14755

Ruach, R., Yellinek, S., Itskovits, E., Deshe, N., Eliezer, Y., Bokman, E., Zaslaver, A., 2022. A negative feedback loop in the GPCR pathway underlies efficient coding of external stimuli. Mol. Syst. Biol. 18, e10514. 10.15252/msb.202110514

Speese, S., Petrie, M., Schuske, K., Ailion, M., Ann, K., Iwasaki, K., Jorgensen, E.M., Martin, T.F.J., 2007. UNC-31 (CAPS) is required for dense-core vesicle but not synaptic vesicle exocytosis in Caenorhabditis elegans. J. Neurosci. Off. J. Soc. Neurosci. 27, 6150–6162. 10.1523/JNEUROSCI.1466-07.2007

Toyoshima, Y., Tokunaga, T., Hirose, O., Kanamori, M., Teramoto, T., Jang, M.S., Kuge, S., Ishihara, T., Yoshida, R., Iino, Y., 2016. Accurate Automatic Detection of Densely Distributed Cell Nuclei in 3D Space. PLOS Comput. Biol. 12, e1004970. 10.1371/journal.pcbi.1004970

Varoqueaux, F., Sigler, A., Rhee, J.-S., Brose, N., Enk, C., Reim, K., Rosenmund, C., 2002. Total arrest of spontaneous and evoked synaptic transmission but normal synaptogenesis in the absence of Munc13-mediated vesicle priming. Proc. Natl. Acad. Sci. 99, 9037–9042. 10.1073/pnas.122623799

White, J.G., Southgate, E., Thomson, J.N., Brenner, S., 1986. The structure of the nervous system of the nematode Caenorhabditis elegans. Philos. Trans. R. Soc. Lond. B. Biol. Sci. 314, 1–340.

Witvliet, D., Mulcahy, B., Mitchell, J.K., Meirovitch, Y., Berger, D.R., Wu, Y., Liu, Y., Koh, W.X., Parvathala, R., Holmyard, D., Schalek, R.L., Shavit, N., Chisholm, A.D., Lichtman, J.W., Samuel, A.D.T., Zhen, M., 2021. Connectomes across development reveal principles of brain maturation. Nature 596, 257–261. 10.1038/s41586-021-03778-8

Zaslaver, A., Liani, I., Shtangel, O., Ginzburg, S., Yee, L., Sternberg, P.W., 2015. Hierarchical sparse coding in the sensory system of Caenorhabditis elegans. Proc. Natl. Acad. Sci. 112, 1185–1189. 10.1073/pnas.1423656112

Zhou, K., Stawicki, T.M., Goncharov, A., Jin, Y., 2013. Position of UNC-13 in the active zone regulates synaptic vesicle release probability and release kinetics. eLife 2, e01180. 10.7554/eLife.01180

